# *bioSyntax*: Syntax Highlighting For Computational Biology

**DOI:** 10.1101/235820

**Authors:** Artem Babaian, Anicet Ebou, Alyssa Fegen, Ho Yin (Jeffrey) Kam, German E. Novakovsky, Jasper Wong

## Abstract

Computational biology requires the reading and comprehension of biological data files. Plain-text formats such as SAM, VCF, GTF, PDB and FASTA, often contain critical information that is obfuscated by the complexity of the data structures. *bioSyntax* (<u>http://bioSyntax.org</u>) is a freely available suite of syntax highlighting packages for *vim, gedit, Sublime*, and *less*, which aids computational scientists to parse and work with their data more efficiently.

## Background

A major component of computational biology research involves the reading and comprehension of data in biological file-formats including FASTA[1], FASTQ[2], gene transfer format (GTF)[3], variant calling format (VCF)[4], protein database format (PDB) [5,6] and, sequence alignment map (SAM)[7] amongst others[8]. While being easy to parse computationally, these and other biological files often become illegible to scientists as their size and complexity increases, including header sections containing critical data descriptors often required for downstream processing.

Syntax highlighting is designed to improve the interpretability of text files with well-defined structures through the application of colour, font, and formatting to differentiate content; typically a set of keywords, structures, and symbols. Originally developed for the code editor Emily in 1971[9] and later LEXX in late 1980s[10], syntax highlighting is now ubiquitous in the computer sciences. Syntax highlighting reduces task completion times for information processing compared to plain text, thereby improving content comprehension[11,12].

A plethora of tools, both stand-alone and web-based, exist to help scientists visualize and process biological data[13–19]. However, these applications require extra usertime to learn and competently operate, and do not load as quickly as a simple text editor or pager.

The objective of *bioSyntax* is to improve the human readability of scientific data-formats through seamlessly integrated syntax highlighting and to assist scientists in working with low-level data files. *bioSyntax* is currently ported for three common text editors, *Vim* (and *GVim*), *gedit* (and other linux editors using the *GTKSourceView* library) and *Sublime-Text-3* as well as the command-line pager program, *less*. Additionally, *bioSyntax* functions as a repository into which syntax highlighting definition files may be deposited by the community for additional scientific file formats.

## Results

### Syntax Highlighting for Computational Biology File Formats

*bioSyntax* currently recognizes FASTA, FASTQ, CLUSTAL, BED, GTF, PDB, SAM and VCF formats across all three text editors and *less*. Upon installation, *bioSyntax* automatically recognizes file-extensions and seamlessly assign syntax highlighting to these data files.

The main benefit of syntax highlighting is immediately apparent through its increased legibility (Figure 1, Supplementary Figures 1 and 2), especially in the deconstruction of verbose content such as plain-text CIGAR strings (Figure 1A). Each file-format uses contrasting colours to accentuate keywords or key fields. Nucleotides and amino acids are highlighted with distinct colours allowing for users to read sequences and interpret patterns in the alignment. Data fields containing scores such as PHRED base quality or mapping scores are gradient coloured.

**Figure 1:**
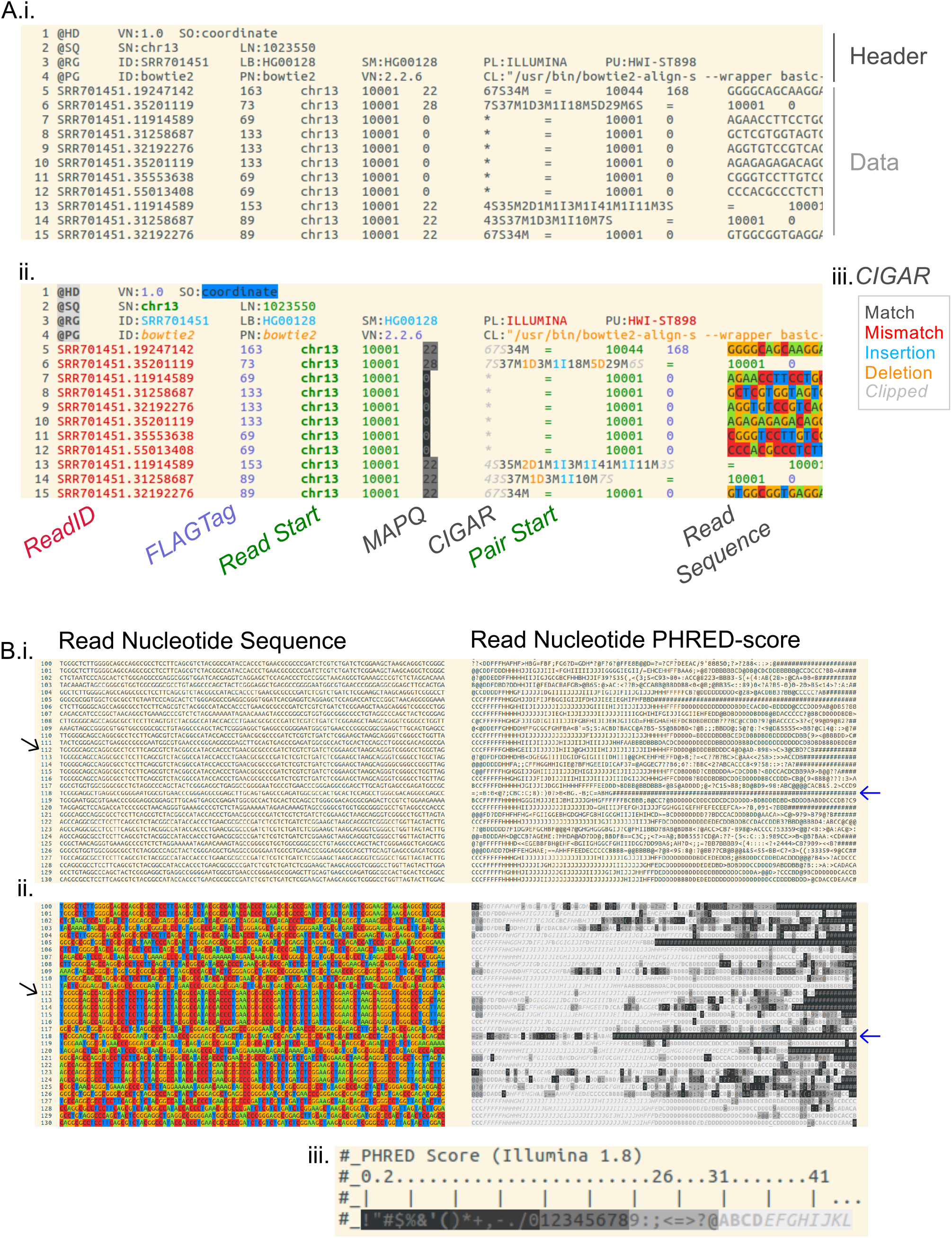
Syntax highlighting for sequence alignment map (.sam) file format. **A)** Terminal screenshot of the `‘less HG00128_hgr1.sam’` command run i. normally or ii. with bioSyntax. Related information in the header and data sections are grouped by colours (genomic coordinates, green; sample information, pale blue …) to improve legibility. Each data-row is an individual sequencing read. iii. CIGAR alignment strings in particular can be highlighted such that they become substantially easier to read. **B)** A broad view of the nucleotide and PHRED-score for 30 reads i. before, and ii. after syntax highlighting. Underlying information of about the data becomes intuitively visible such as PCR-duplicates (black arrow) and poor quality areas and reads (blue arrow) based on iii. PHRED score.

The overall system of highlighting is also designed to group biological classes, even across file formats (Supplementary Table 1). For instance, dark-green is reserved for genomic coordinates in BED, GTF, SAM and VCF, so even if a user is unfamiliar with the SAM format, previous experience associating dark-green in BED or GTF will inform them of the meaning of those fields when presented in a SAM file (Supplementary Figure 2).

Ultimately, *bioSyntax* aims to help computational biologists comprehend data using intuitive graphical highlighting rather than simple syntax highlighting. When the data does not have to be read per character or per word, but can be viewed as patterns, underlying information in the data becomes apparent, similar to alternative nucleotide representations[20,21]. This is best seen in complex files such as SAM in which PCR-duplicate reads form block patterns and read density can be approximated by the diagonal similarity of reads at a locus (Figure 1B).

### bioSyntax nucleotide representation

*bioSyntax* implements a novel nucleotide colouring scheme for the complete IUPAC ambiguous base set[22], unlike other colour-sets which are designed for four or five bases (Figures 2). The four primary base colours are chosen such that additive colour mixing also represents complete base ambiguity. For instance, thymine (blue) and cytosine (red) are pyrimidines (magenta), and the any base, N, is white. This colour-set visually distinguishes the strong bases (G,C) and weak bases (A,T) as warm and cool colours respectively, allowing for an intuitive approximation of a sequence GC-content (Figure 2C). Additionally, a high-contrast colour-scheme is available to aid visually impaired or colour-blind users (Supplementary Figure 3).

**Figure 2:**
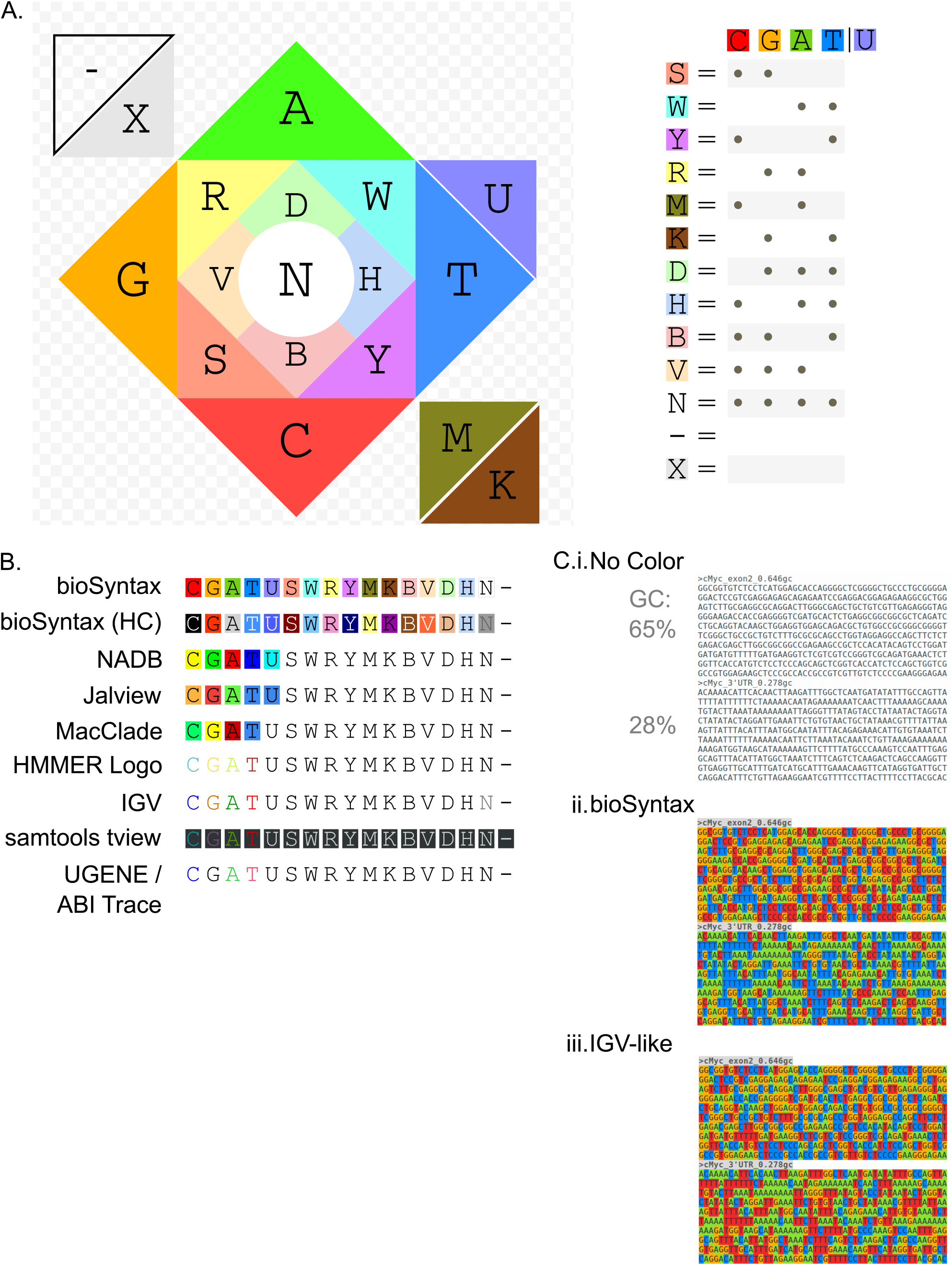
*bioSyntax* nucleotide colour scheme. **A)** The four primary bases are coloured in two pairs of contrasting colours. IUPAC ambiguous bases are then coloured in increasingly lighter tones of the approximately mixed colours. To accomodate 4-dimensional bases in 3-dimensional colours, aMino (A or C) and Keto (G or T) bases are darker. **B)** A comparison of nucleotide colour-schemes in the literature. C) bioSyntax colouring allows for approximation of a sequences GC-content by how warm (high GC) or cool (high AT) it appears.

### The bioSyntax repository

There are scores of biological and scientific file-formats which would benefit from syntax highlighting. To facilitate future development of syntax definition files in science, the *bioSyntax* repository (<u>http://bioSyntax.org/repo</u>) was set-up. The repository is both a library for scientific syntax highlighting and a community-oriented resource for learning syntax highlighting development. In this manner, researchers experienced in the use-cases of a file-format can quickly develop and share new syntax definition files.

## Conclusions

*bioSyntax* allows researchers to intuitively read and navigate biological files in the context of familiar and common text editing tools. The cross-format unifying colour theme for biological data classes aids users in the rapid and accurate parsing of data, even with minimal prior knowledge regarding the file-format. Altogether, bioSyntax substantially improves the legibility of biological data and helps researchers to grok their data.

## Methods

### Availability and Use

*bioSyntax* installers are freely available at <u>http://bioSyntax.org</u> for *vim* (Linux, MacOS, Windows), *less* (Linux, MacOS), *gedit* (Linux, MacOS, Windows) and *sublime-text-3* (Linux, MacOS, Windows) under GNU General Public License v3.0. Source code is available at <u>http://www.github.com/bioSyntax/bioSyntax</u>.

*bioSyntax* for *less* depends on the *source-highlight* package. For *gedit* syntax highlighting, *bioSyntax* was developed for the requisite *GTKSourceView3* library.

Highlighting of FASTA, FASTQ, CLUSTAL, BED, GTF, PDB, VCF and SAM files is automatically detected by file extension and can also be manually set within text-editors for files with non-standard file-extensions. In *gedit, sublime* and *vim*, amino acid FASTA files can be colored using CLUSTAL[23], Taylor[24], Zappo[13] or Hydrophobicity[14] color schemes.

Large and compressed data can be piped directly into *less* using the commands: `‘sam-less’, ‘vcf-less’`, etc…(Figure 1). For example *‘samtools view −h NAl2878.bam* | *sam-less - ’*, or *‘gzip −dc gencode.v26.gtf.gz* | *grep ‘MYC’*; - | *gtf-less −x 10* - ’.

### File Specifications

At its core, bioSyntax is a set of syntax-highlighting definition files which are themselves a programmed set of regular expressions.

Where available, syntax-files are designed using a combination of; official file specifications SAM v1.5 (<u>https://samtools.github.io/hts-specs/SAMv1.pdf</u>), VCF v4.2 (<u>https://samtools.github.io/hts-specs/VCFv4.2.pdf</u>) PDB v3.30 (<u>ftp://ftp.wwpdb.org/pub/pdb/doc/format_descriptions/Format_v33_Letter.pdf</u>), BED6 (<u>https://genome.ucsc.edu/FAQ/FAQformat.html</u>), GTF v2.2 (<u>http://mblab.wustl.edu/GTF22.html</u>), example files from databases (NCBI Nucleotide/Protein[25], Sequence Read Archive[26], dbSNP[27], RefSeq[28], RCSB[5] and UCSC Genome Browser[17]); publically available consortium data (1000 genomes project[29], ENCODE[30]); and standard outputs from commonly used software (*samtools[7], GATK[31], bowtie2[32], cufflinks[33]* and *Clustalx[23]*).

## List of Abbreviations

GTF: : Gene Transfer Format
IUPAC: : International Union of Pure and Applied Chemistry
PDB: : Protein Database Format
SAM: : Sequence Alignment Map
VCF: : Variant Call Format

## Supplementary Figure and Table Legends

**Supplementary Figure 1:**
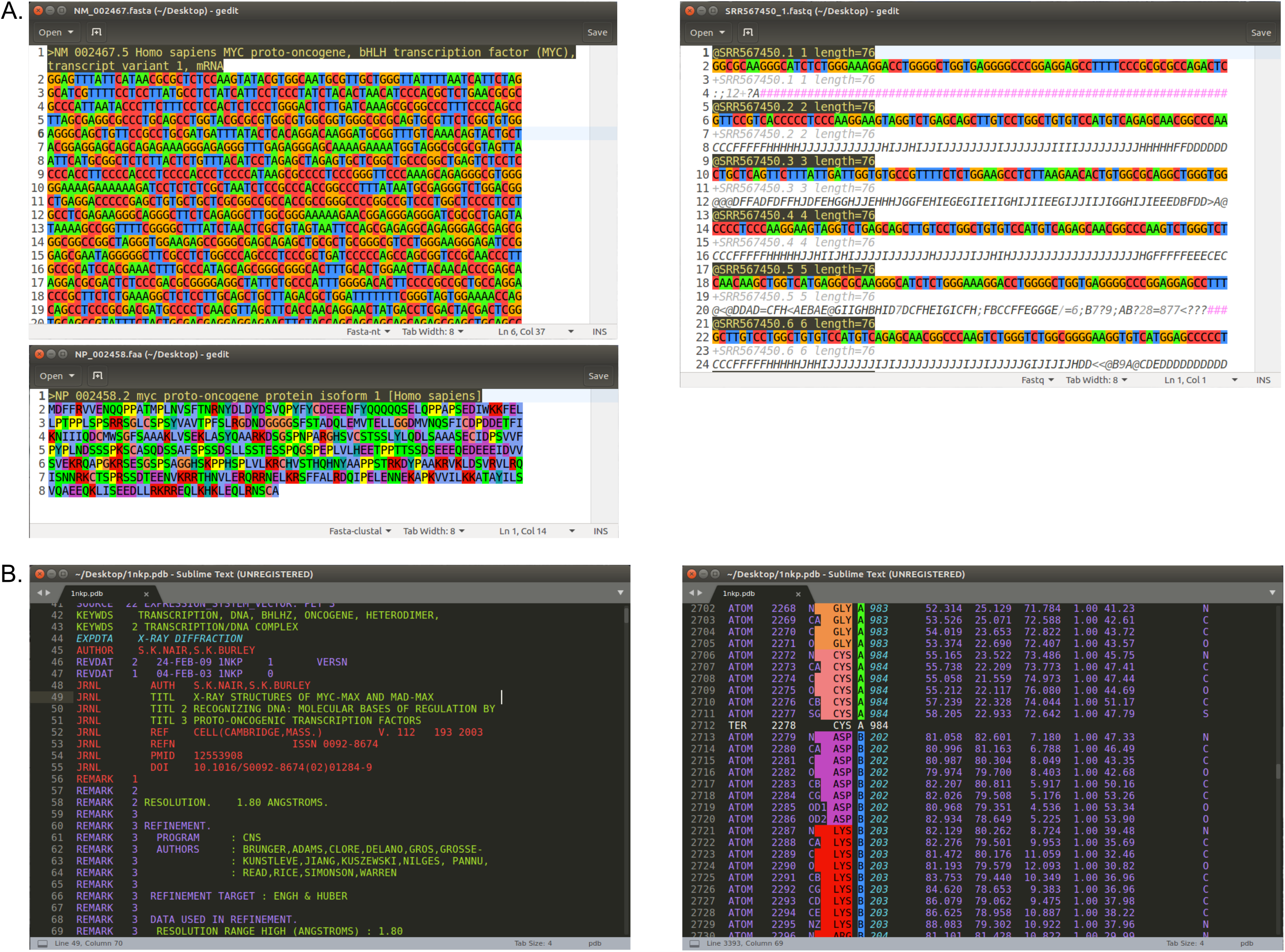
Screenshots of *bioSyntax* in; **A)** *gedit* for the nucleotide sequence FASTA, an amino acid FASTA with CLUSTAL color scheme and a FASTQ file and; **B)** *sublime-text-3* for a PDB file.

**Supplementary Figure 2:**
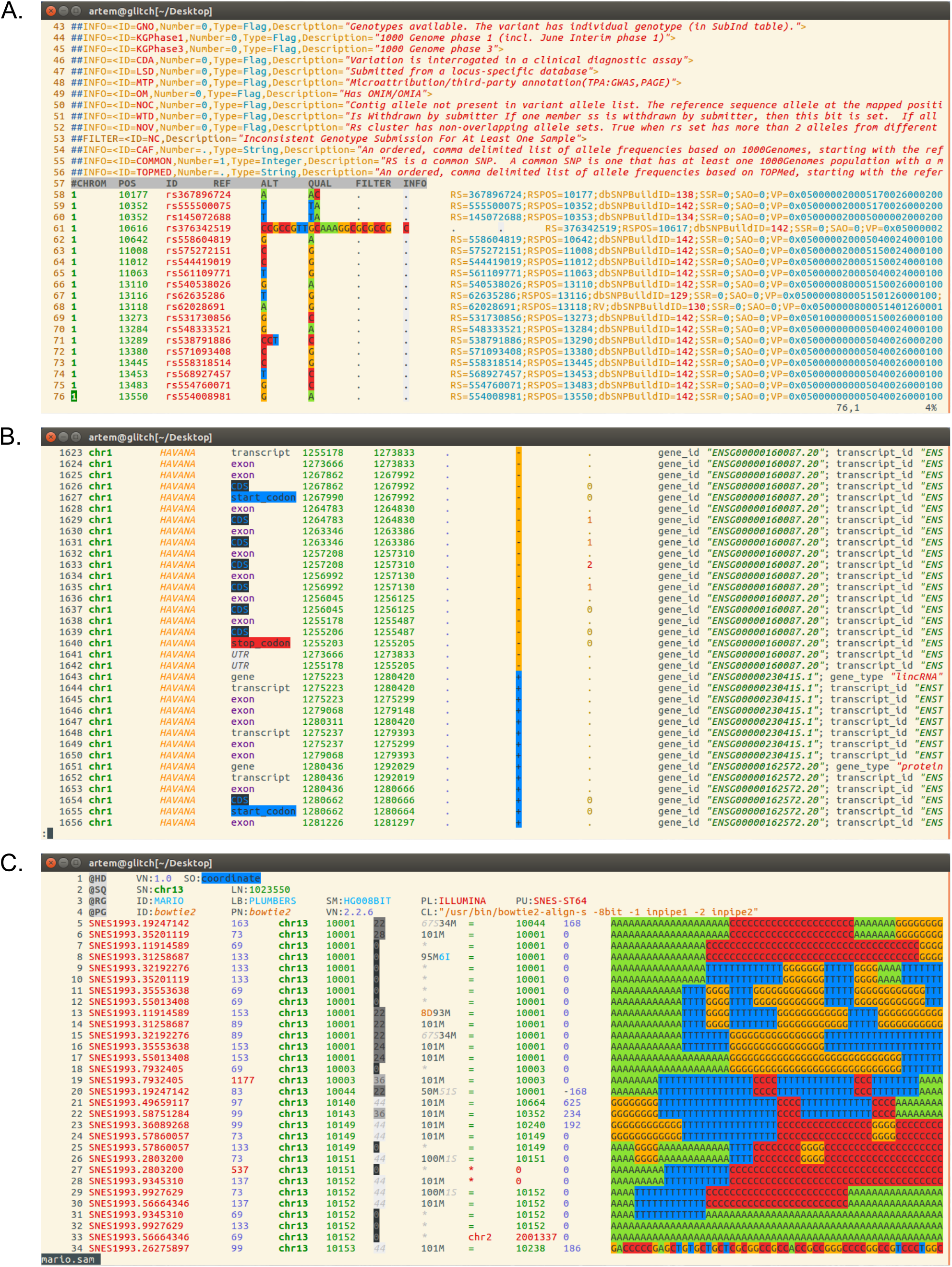
Screenshots of *bioSyntax* in; **A)** *vim* for the human dbSNP (hg38 build-150) VCF file and; **B)** *less* for the Gencode v26 Annotation GTF and **C)** an example SAM file. In the GTF format, note how background colouring of “start_codon”, “stop_codon”, “CDS”, and “UTR” graphically distinguishes protein-coding transcripts from non-coding transcripts.

**Supplementary Figure 3:**
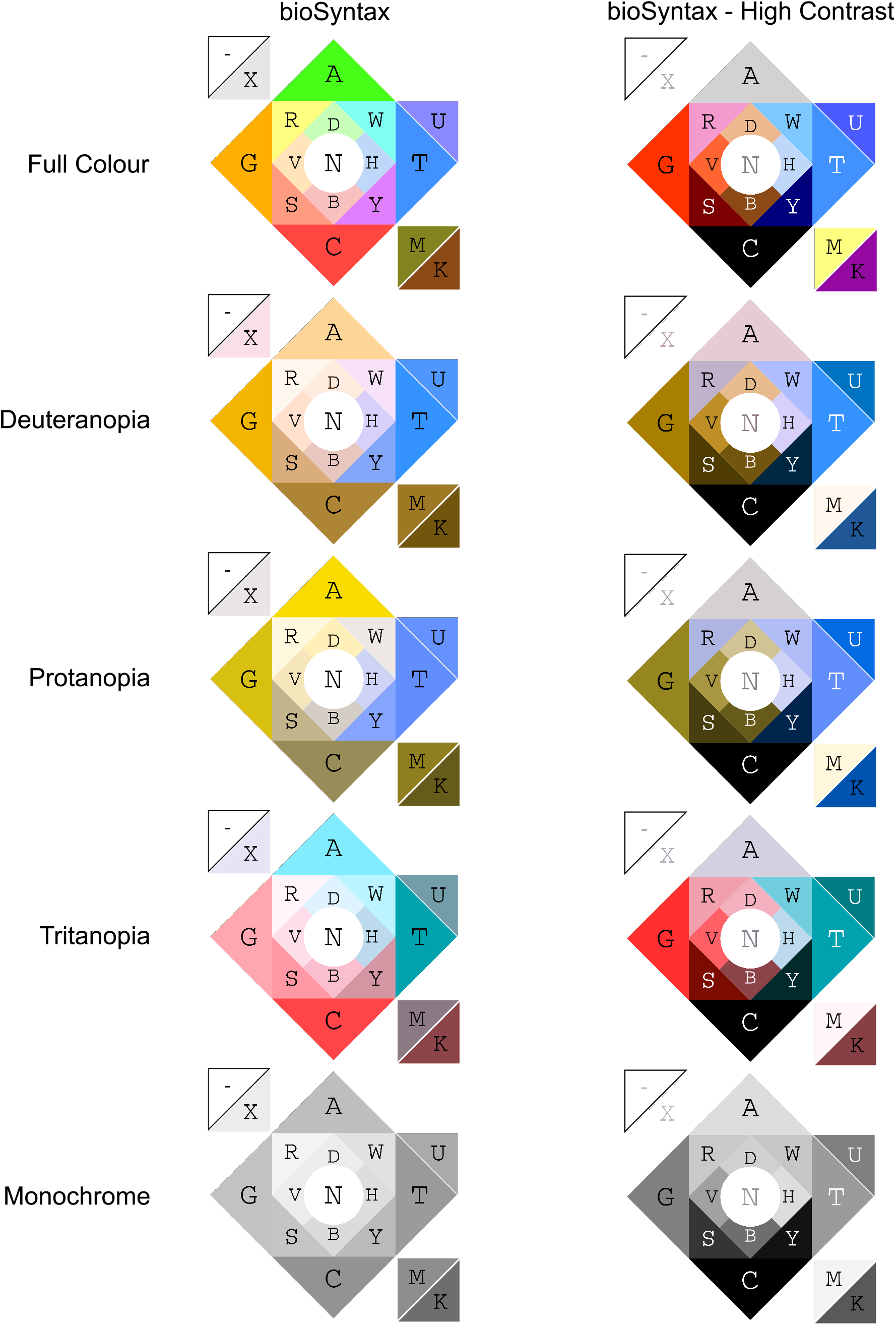
The standard and high-contrast bioSyntax color set for IUPAC nucleotides under the different forms of simulated color-blindness. Hue and lightness variations in the standard theme allow for accesibility with color-blindness. The alternative high-contrast set retains higher visual distinction between bases, even at the monochrome level.

*Supplementary Table 1:*

Biological class definitions and colour definitions for the default bioSyntax theme in hexadecimal (*gedit, sublime, gvim*), cterm (*vim*), and 8-bit ANSI escape character (*less*) colours.

## Author Contributions

AB conceived and led the design/development of *bioSyntax*. All authors contributed to coding and implementation. All authors wrote, read and approved the final manuscript.

## Acknowledgements

*bioSyntax* was initiated at the *hackseq17* (<u>http://www.hackseq.com</u>) genomics hackathon[34]. We would like to thank the organizers for their dedication to making the event happen and ongoing support during development. We’d like to thank Ching Pan Eric Chu for early work on developing regular expressions for the SAM format and Joseph O’Brien for his artistic insight in developing a high-contrast palette for nucleotide colouring.

## Ethics Approval

Not applicable.

